# Visuomotor information drives interference between the hands more than dynamic motor information during bimanual reaching

**DOI:** 10.1101/2021.03.15.435502

**Authors:** Phillip C. Desrochers, Alexander T. Brunfeldt, Florian A. Kagerer

## Abstract

During complex bimanual movements, interference can occur in the form of one hand influencing the action of the contralateral hand. Interference likely results from conflicting sensorimotor information shared between brain regions controlling hand movements via neural crosstalk. However, how visual and force-related feedback processes interact with each other during bimanual reaching is not well understood. In this study, four groups experienced either a visuomotor perturbation, dynamic perturbation, combined visuomotor and dynamic perturbation, or no perturbation in their right hand during bimanual reaches, with each hand controlling its own cursor. The left hand was examined for interference as a consequence of the right-hand perturbation. The results indicated that the visuomotor and combined perturbations showed greater interference in the left hand than the dynamic perturbation, but that the combined and visuomotor perturbations were equivalent. This suggests that dynamic sensorimotor and visuomotor processes do not interact between hemisphere-hand systems, and that primarily visuomotor processes lead to interference between the hands.

## Introduction

For humans, bimanual actions comprise a significant part of everyday life. While the adult neuromotor system is usually capable of performing simple or well-practiced bimanual tasks without problems (e.g., clapping, tying one’s shoes, typing), more complicated movements, particularly those with spatiotemporal incongruencies, can produce interference between the hands (e.g., the familiar childhood trick of rubbing one’s belly while patting one’s head, juggling, or playing the drums; Franz *et al*., 1991). Interference is a phenomenon where the action of one hand can influence the action of the other hand. Interference during bimanual coordination has been well studied (Semjen *et al*., 1995; Kennerley *et al*., 2002; Hazeltine *et al*., 2003; Obhi & Goodale, 2005; Kagerer, 2016; Shea *et al*., 2016), and is thought to result from neural crosstalk between brain areas controlling bimanual movements. However, the specific type of sensorimotor information (i.e., direction, force etc.) being shared between the hemispheres, and how this information might interact in the context of interference, remains an open question.

It is possible to probe the function of different types of sensorimotor information within the CNS by asking participants to adapt to perturbations which manipulate specific sensorimotor components. To probe visuomotor processes, the visual feedback received by a participant can be manipulated to introduce a discrepancy between the intended and observed movement consequences. To probe dynamic motor processes, the forces experienced during a movement can be manipulated to likewise provide unexpected dynamic feedback. Such perturbations require participants to modify so-called internal models (i.e., neural representations of the external world) and sensorimotor feedback gains to minimize the movement error introduced by the perturbation (Wolpert *et al*., 1995; Kawato, 1999; Körding & Wolpert, 2004; Izawa & Shadmehr, 2008; Franklin *et al*., 2012). When these perturbations are applied to one hand during bimanual reaches, spatial interference in the opposing hand can be inferred to be due to sharing of motor information and internal representations between hemisphere-hand systems in that specific sensorimotor domain (Kagerer, 2015a).

It has been shown previously that visuomotor perturbation of the right hand causes interference in the left hand (Kagerer, 2015b, 2016; Desrochers *et al*., 2020). However, recent evidence demonstrated that a dynamic perturbation does not produce substantial interference (Desrochers et al., 2017). While surprising, given that both perturbations theoretically require the updating of internal models, this finding is supported by research showing that learning of a dynamic perturbation in one hand neither interferes with nor facilitates learning of a separate dynamic perturbation in the opposing hand (Tcheang *et al*., 2007). Furthermore, different neural processes may be involved in the adaptation to visuomotor vs. dynamic perturbations, since one can be learned without simultaneously interfering with the other (Krakauer *et al*., 1999), though this does not necessarily hold when visuomotor and dynamic perturbation act with respect to the same kinematic variable (Tong *et al*., 2002). Several studies suggest that visual information dominates trajectory control over proprioceptive information (Judkins & Scheidt, 2013; Sexton *et al*., 2019). For example, when participants are not provided with visual errors orthogonal to the direction of reaching, adaptation to a dynamic perturbation suffers, suggesting that visuomotor and dynamic motor information compete or regulate different phases of movement (Scheidt *et al*., 2005). The dominant sensory modality affecting a reach may also depend on the sensory information that informs the initial hand position, and thus, the motor plan (Sarlegna & Sainburg, 2007). Fewer studies have investigated interplay between visual and dynamic sensory information in the context of bimanual coordination. Swinnen et al. (2001) demonstrated that adding force constraints to a bimanual spatial interference task did not result in additional interference. Finally, Diedrichsen (2007) showed that when participants control individual cursors with each hand and reach to separate targets, a force perturbation in the right hand did not influence the trajectory of the left hand. Thus, in this study, we specifically assessed whether visuomotor and dynamic perturbations of similar magnitude contribute differently to interference between the hands.

It appears, however, that dynamics are still pertinent to motor learning and interference paradigms. Unimanual studies have shown an interaction between visuomotor and dynamic perturbations within the sensorimotor system. Franklin and colleagues (2012) demonstrated that while adapting to a dynamic perturbation, participants’ motor responses to a visuomotor perturbation are upregulated compared to when they are not adapting to a dynamic perturbation. These findings suggest that the presence of task relevant dynamics may increase sensorimotor gain and upregulate responses to visuomotor perturbations while feed-forward internal models are being updated. In other words, adaptation to one type of sensorimotor information may increase responses to perturbations of the other type. However, it is unknown whether, during a bimanual task, upregulating sensorimotor networks via simultaneous dynamic and visuomotor perturbations in one hemisphere-hand system will influence the contralateral hemisphere-hand system, resulting in increased interference compared to either perturbation alone.

In this experiment, we examined how visuomotor, dynamic, and combined visuomotor and dynamic perturbations in the right hand affected interference in the left, unperturbed hand. We hypothesized that if visuomotor and dynamic motor information are differentially shared between hemispheres, interference due to perturbations of similar magnitude will be observed at disparate levels. Additionally, if the motor system responds to dynamic and visuomotor perturbations synergistically, as suggested by Franklin and colleagues (2012), interference in the left hand due to a simultaneous dynamic and visuomotor perturbation of the right hand would be greater than interference during dynamic or visuomotor perturbation alone.

## Methods

### Participants

We recruited sixty right-handed young adults, confirmed via the Edinburgh Handedness Inventory – Short form (Oldfield, 1971), who were between 18 and 30 years old. Participants had no history of cognitive or neurological impairment, had not sustained a concussion in the past year, and had normal or corrected-to-normal vision. All participants provided informed consent, and all procedures were approved by the Michigan State University Institutional Review Board.

### Study design

Participants were randomly assigned to one of four groups (15 participants per group; Figure 1): no perturbation, visuomotor perturbation, dynamic perturbation, or combined perturbation (simultaneous dynamic + visuomotor; Figure 1). Participants were cued to move two Kinarm End-Point Lab (BKIN Technologies Ltd., Kingston, Ontario) manipulanda simultaneously from two home positions to two targets located 10 cm directly forward (90°) or backward (270°) from the home positions. They were instructed to reach straight, fast, and accurately from the home position to the target. Targets were randomly presented, but both hands always moved in the same direction. After holding steady in the target position for 1500 ms, participants were cued to return to the home positions. Hand position was represented by a cursor on a screen that occluded vision of the hands. Participants started by performing unperturbed reaches during two blocks of 30 trials. In the first “visual baseline” (VBL) block, hand feedback was displayed for both hands. Then, in the “kinesthetic baseline” (KBL) block, visual feedback (cursor position) was removed for the left hand, requiring participants to rely on kinesthetic control. Participants were instructed to continue reaching to the target with their nonvisible hand, stopping when they estimated that their hand was in the target. Then, in the “exposure” (EXP) block of 250 trials, participants were exposed to the perturbation (Figure 2). For the visuomotor perturbation group, the cursor representing the right hand was rotated 45° clockwise about the home position, such that participants needed to adapt their right-hand movement trajectory −45° to hit the target. For the dynamic perturbation group, the participants encountered a curl field force acting 90° perpendicular to the instantaneous velocity vector with a magnitude of 20 N per m/s of the reach velocity magnitude. The combined perturbation group received both perturbations simultaneously, such that the visual feedback was rotated 45° and a 20 N per m/s force perpendicular to the direction of instantaneous velocity was applied. The control group experienced no perturbation in the right hand. Finally, in the “post-exposure” (Post-EXP) block of 50 trials, the perturbations for the three experimental groups were removed. During the EXP and Post-EXP blocks, left hand visual feedback remained off, leaving the left hand susceptible to interference.

**Figure 1:**
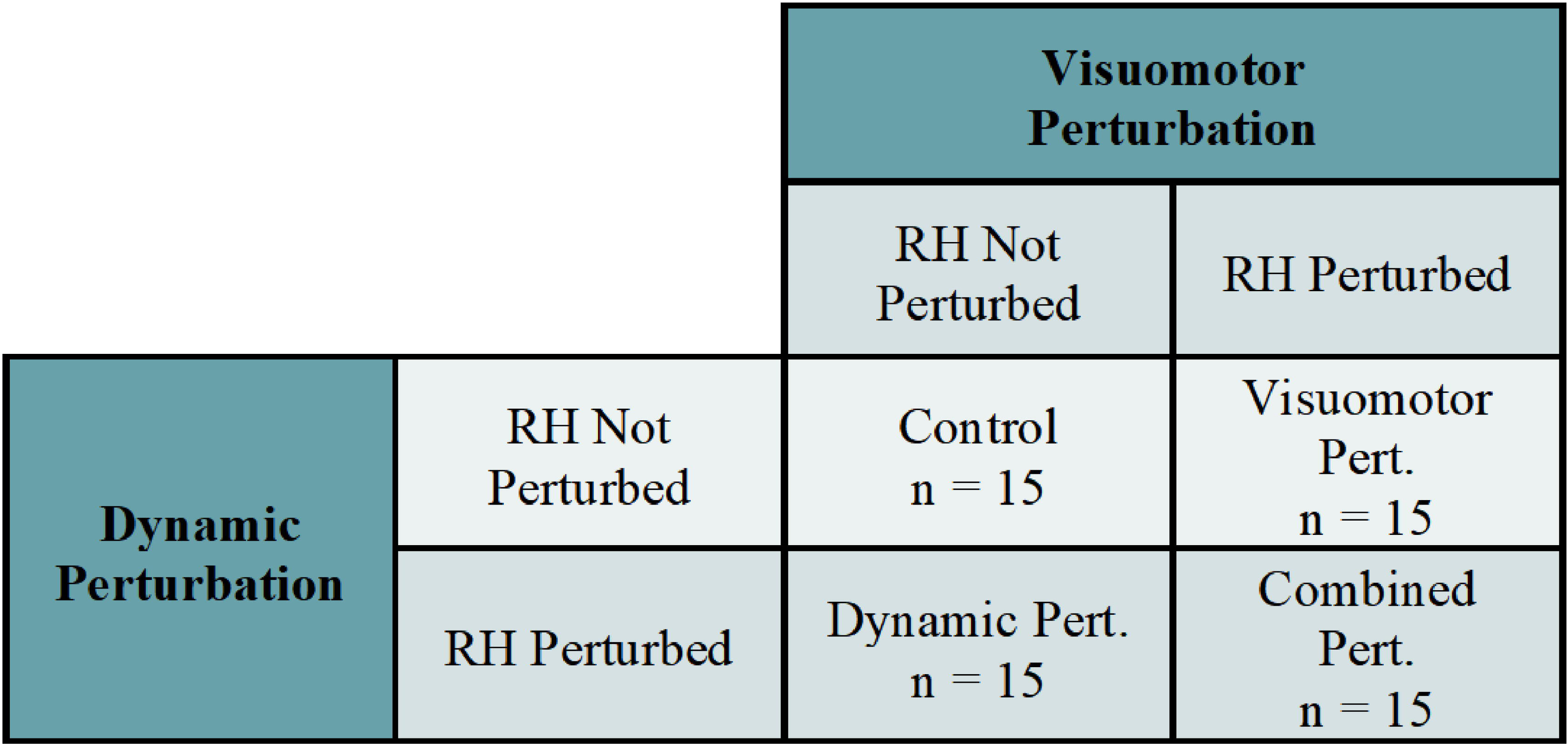
Participants received no perturbation (control), a 40° visual rotation (visuomotor perturbation), a 20 Nsm-1 velocity dependent force (dynamic perturbation), or both perturbations (combined perturbation) in their right hand.

**Figure 2:**
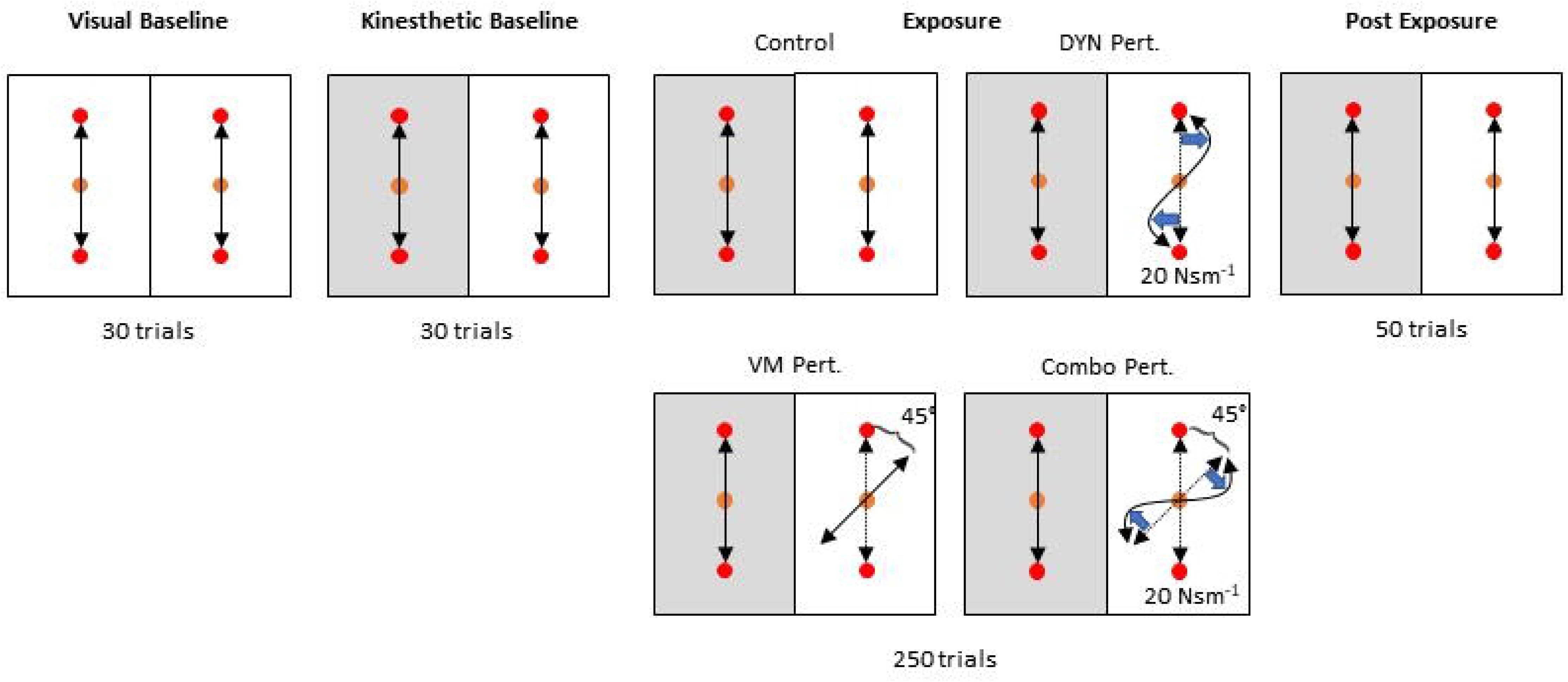
Boxes show the task design for the visual baseline, kinesthetic baseline, exposure, and post-exposure blocks. In the exposure blocks, the different groups received null (control), visuomotor, dynamic, or combined visuomotor and dynamic perturbations. Orange circles represent the home position, and red circles represent the targets. Shaded boxes indicate lack of visual feedback for the corresponding hand.

To ensure that all participants achieved the similar reaching velocities (since the dynamic perturbation is velocity dependent), participants were given feedback regarding the time elapsed between initiation of the reach and the moment both hands were in the targets and had a between-hand mean velocity under 3 cm/s. If that time was under 300 ms, a red rectangle was displayed in the center of the screen, indicating that the reach duration was too short; if the time was over 450 ms, a blue rectangle was displayed (reach duration too long), and if the time was within the 300-450 ms window, a green rectangle was displayed (ideal reach duration). Participants were generally quite good at adhering to the timing constraints, particularly during unperturbed trials and at the end of the exposure period. Trials in which participants did not successfully hit the timing window were not excluded from analysis; subjects were simply reminded to continue trying to achieve movements with the proper velocity.

### Kinematic and kinetic outcome measures

Movement onset and offset were semi-automatically identified using custom written Matlab scripts (Mathworks, inc., Natick, MA) with an onset/offset algorithm (Teasdale *et al*., 1993) and were subsequently checked for accuracy by trained experimenters. To evaluate motor performance in the right hand and interference in the left hand, we assessed kinematic error via four primary outcome measures at each trial: 1) root mean square error (RMSE), a measure of movement straightness across the entirety of the reach, was calculated with reference to a straight line between the targets and normalized to movement length; 2) Initial directional error (IDE), a measure of feed-forward or predictive error of the motor plan, was computed at the moment of peak tangential velocity as the angle between a vector from the home position to the position of the cursor and a vector from the home position to the target; 3) Initial endpoint error (IEE), computed at end of the initial ballistic movement, or the moment where the primary reach ends (i.e., before any subsequent corrective movements) as the angular deviation from the vector between the home position and the hand and the vector between the home position and the target. The end of the initial ballistic movement was defined as the first moment after peak tangential velocity where the hand’s velocity experienced a local minimum (i.e., the start of a secondary movement) or where the hand’s velocity crossed zero (i.e., a reversal of direction). IEE represents a moment from which on afferent sensory information was beginning to be integrated into the sensorimotor system, and online error correction was beginning to occur (Seidler, 2006); 4) Final endpoint error (FEE), computed for the left hand as the lateral displacement between the hand position and the target at the moment when all movement had ceased in that hand. FEE occurred at a time in which the sensorimotor system had integrated all relevant feedback and corrected any perceived movement error.

We also applied a virtual force channel to the left hand in 20% of trials that constrained movement to a straight-line path to the target. The width of the channel was 0.25 mm, while the channel walls were 2 cm thick. The robots linearly applied 20 N/cm as participants moved laterally into the walls. A 5 N lateral viscous force dampened possible harmonic oscillations between the two walls. The channels turned on gradually over a 1500 ms interval after the participants returned to and stopped in the home positions following the previous trial. When participants were informally asked after the experiment whether they noticed any manipulations in the left hand apart from loss of visual feedback, they did not report noticing the force channel. Force sensors embedded in the Kinarm manipulanda endpoints measured the lateral force applied against the ‘channel walls’ and served as an indicator of interference in the left hand. Each reach was interpolated to 1000 data points. The movement trajectories were averaged within early-EXP (first 30 trials of exposure) and late-EXP (last 30 trials of exposure). The force applied to the end-point handles was assessed at three timepoints during the reach: the average point of peak velocity (where IDE was calculated), the average point of the end of the initial ballistic movement (where IEE was calculated), and at the end of the movement (where FEE was calculated).

### Data processing and statistical analyses

Outcome measures for each dependent variable were baseline corrected to the KBL block by subtracting the KBL block mean from each trial before averaging. Kinematic and kinetic measures were evaluated in blocks that comprised the first and last 30 trials in the EXP period. A 2 (Block: Early, Late) x 4 (Group: Control, Visuomotor Perturbation, Dynamic Perturbation, Combined Perturbation) mixed-design ANOVA was used to assess adaptation in the right hand and interference in the left hand for each dependent measure. If violations of sphericity were present, they were adjusted with a Huynh-Feldt correction. Effect sizes were computed using generalized eta squared (*η_g_^2^*; (Bakeman, 2005)). Subsequent post-hoc analyses were performed by collapsing across non-significant independent variables, or by examining simple effects within blocks using one-way ANOVAs when significant main effects or interactions were present. Tukey HSD was used to determine differences between groups within each block. Statistical analyses were performed using custom written scripts in RStudio 1.2 software (RStudio Inc., Boston, MA) using the ‘afex’, ‘ez’, and base R packages.

## Results

### Kinematic measures

First, we wanted to confirm that the magnitudes of the visuomotor and dynamic perturbations of the right hand were comparable. Cursor trajectories are plotted in Figure 3 across Early-EXP and Late-EXP for both hands. An independent samples t-test was used to evaluate the kinematic error in the right hand in the first 10 trials of EXP in the visuomotor and dynamic perturbation groups, before any substantial adaptation took place. No significant difference between the visuomotor and dynamic perturbation groups was found for RMSE (*t*(28) = −0.65, *p* = 0.52) or IEE (*t*(28) = 0.22, *p* = 0.83), though IDE was significant (*t*(28) = 7.98, *p* < 0.001). This showed that while the spatial magnitude of the perturbations was not equivalent at peak velocity, the net effect of the perturbation throughout the full movement trajectory was similar (as shown through IEE and RMSE). Thus, differences between the movements in the early, feed-forward components of the reach in the earliest moments of the EXP block would be expected, even when the overall effects were similar in magnitude.

**Figure 3:**
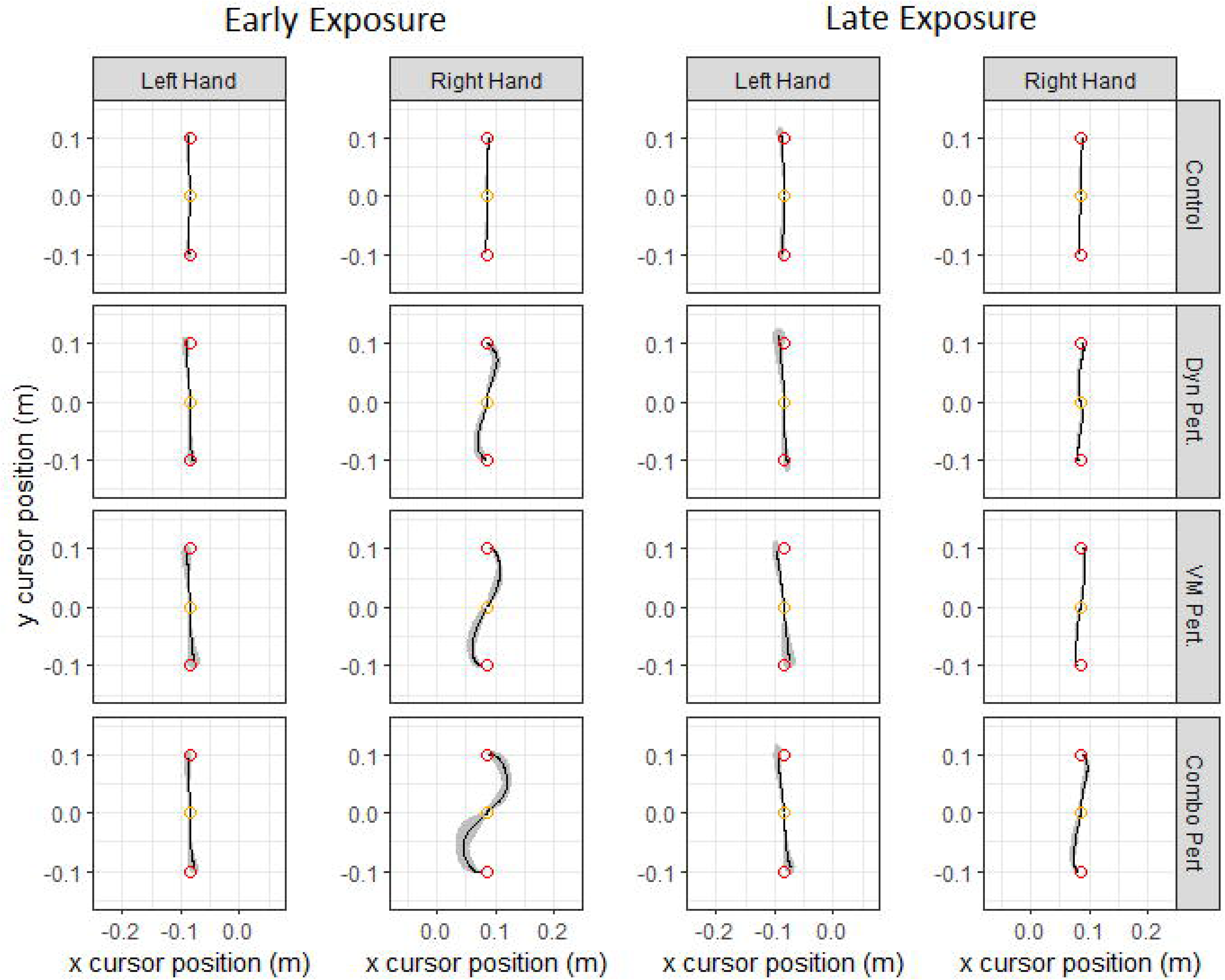
Hand paths for the left and right hands for each group across the Early- and Late-EXP blocks. Lines represent group means, clouds represent standard deviation of group hand paths. Colored circles represent the home position (orange) and targets (red).

Next, we evaluated participants’ adaptation to the respective perturbations. For RMSE across the exposure block, there were significant main effects of group (*F*(3, 56) = 85.16, *p* < 0.0001, *η_g_^2^* = 0.77), and block (*F*(1, 56) = 463.50, *p* < 0.0001, *η_g_^2^* = 0.68), with RMSE decreasing across exposure in the perturbation groups as participants adapted to the perturbations. A significant interaction was also present (*F*(3, 56) = 65.35, *p* < 0.0001, *η_g_^2^* = 0.48), indicating adaptation in the perturbation groups while the control group’s error did not change across exposure. Similar patterns of results were also found for IDE and IEDE (all *p* < 0.0001). As such, subsequent analyses of right-hand adaptation focused on RMSE as a measure of error across the full reach. Within each block of RMSE, one-way ANOVAs revealed significant main effects of group at early-EXP (*F*(3, 56) = 84.44, *p* <0.0001, *η_g_^2^* = 0.82) and late-EXP (*F*(3, 56) = 54.36, *p* < 0.01, *η_g_^2^* =0.74). Tukey HSD tests within each block, with family-wise adjustment for four estimates, showed that all three perturbation groups had significantly greater RMSE than controls at early EXP (all *p* < 0.01). The combined perturbation group showed significantly different RMSE than the dynamic and visuomotor perturbation groups (*p* < 0.0001), while the dynamic and visuomotor perturbation groups did not differ from one another. At late EXP, all groups again showed significantly greater RMSE than the control group (*p* < 0.01). Additionally, all perturbation groups were significantly different from one another (*p* ≤ 0.01), with the greatest RMSE in the combined perturbation group, followed by the dynamic perturbation, visuomotor, and control groups.

We then investigated aftereffects by performing a 2 (Late EXP vs. first 30 trials post-EXP) x 4 (Group) mixed-design ANOVA. If participants had adapted to the perturbation, a subsequent increase in error once the perturbation was removed was expected. Unsurprisingly, we found a main effect of group, showing that RMSE was significantly larger in the perturbation groups following removal of the perturbation (*F*(3, 56) = 104.70, *p* < 0.0001, *η_g_^2^* = 0.74), and a main effect of block, with greater RMSE in the post-EXP block (*F*(1, 56) = 134.37, *p* < 0.0001, *η_g_^2^* = 0.54). A significant interaction was also present (*F*(3, 56) = 22.89, *p* < 0.0001, *η_g_^2^* = 0.37). A similar pattern of findings was observed for both IDE and IEE (all *p* < 0.01). Thus, remaining analyses again focused on RMSE. Analysis of simple effects at post-EXP revealed a significant main effect of group (*F*(3, 56) = 68.38, *p* < 0.0001, *η_g_^2^* = 0.79), and subsequent Tukey HSD post-hoc tests showed that all groups were significantly different from each other (*p* = 0.001), with the exception of the visuomotor and combined perturbation groups, which did not differ (*p* = 0.12). The combined perturbation group showed the greatest aftereffects, followed by the visuomotor perturbation group and dynamic perturbation group. These demonstrate that all perturbation groups were able to successfully adapt to the perturbation and had significant aftereffects once the perturbation was removed (Figure 4).

**Figure 4:**
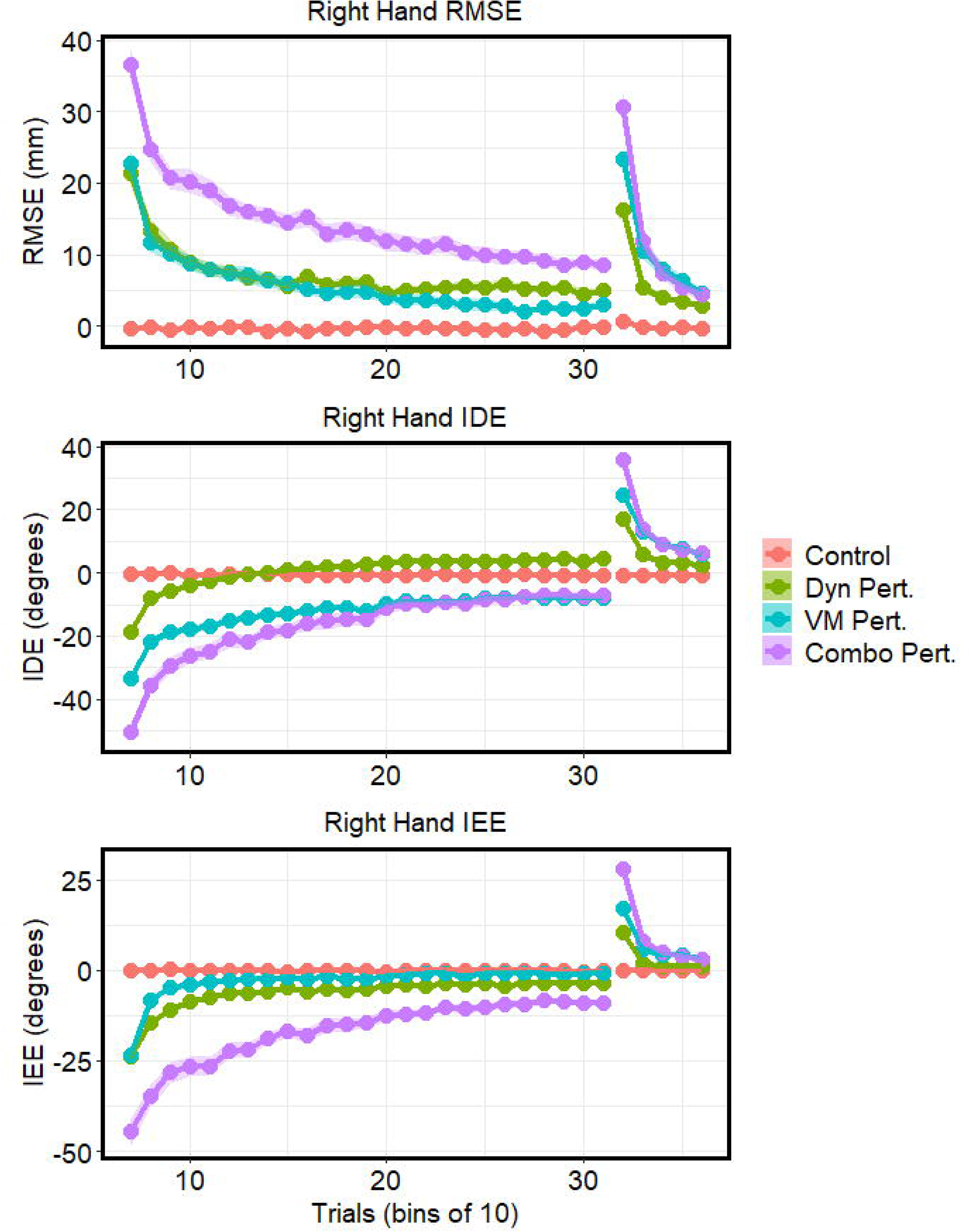
Right hand RMSE, IDE, and IEE across the exposure and post-exposure blocks. Points represent means of 10 consecutive trials. Clouds denote standard error. For IDE and IEE, angles in the first and third quadrants were negative (origin at home position)

After establishing the adaptation characteristics of the groups’ perturbed right hands, we investigated the reaching behavior of the left, interfered-with hand. For RMSE, the overall measure of left-hand reaching error, a mixed-design ANOVA revealed significant main effects of group (*F*(3, 56) = 18.33, *p* < 0.0001, *η_g_^2^* = 0.42) and block (*F*(1, 56) = 25.09, *p* < 0.0001, *η_g_^2^* = 0.11), with RMSE decreasing across the exposure period (Figure 5, A & B), and a significant Group x Block interaction (*F*(3, 56) = 3.62, *p* = 0.02, *η_g_^2^* = 0.05), showing that while interference in left hand changed across exposure for the perturbation groups, movement error remained relatively unchanged for the control group. Examination of the simple effects in both early- and late-EXP showed significant group main effects (all *p* < 0.0001, *η_g_^2^* > 0.30). In early-EXP, Tukey HSD showed that all perturbation groups were significantly different from the controls (all *p* < 0.001). However, after correcting for multiple comparisons, RMSE did not significantly differ between the perturbation groups, though there was a trend for a difference between the combined perturbation and dynamic perturbation groups (*p* = 0.08). In late-EXP, the combined and visuomotor perturbation groups showed significantly higher RMSE than the controls (*p* < 0.01), whereas the dynamic perturbation group did not. Between perturbation groups, the difference between the combined and dynamic perturbation groups was significant (*p* = 0.03). Taken together, this shows that over the whole of the reach, all three perturbation groups showed more interference than controls, with progressively more interference across the dynamic, visuomotor, and combined perturbations.

**Figure 5:**
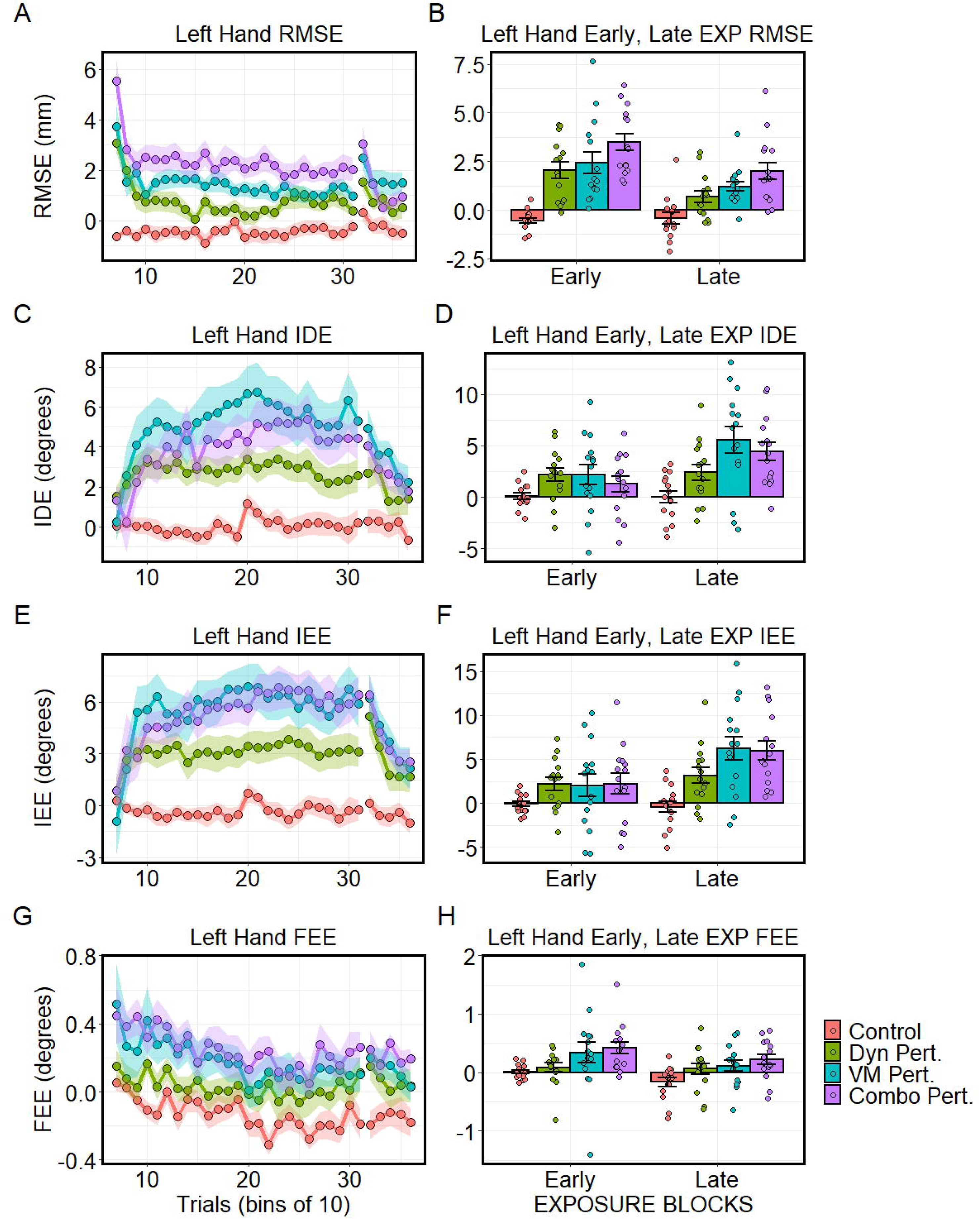
Left hand kinematic dependent variables. A: RMSE, B: IDE, C: IEE, & D: FEE. Means are baseline corrected to the KBL block. For IDE and IEE, angles in the first and third quadrants were negative (origin at home position); thus, interference manifested as a counterclockwise bias in the movement direction. Left panels depict each respective measure in bins of 10 trials each across the exposure (trial bins 7-31) and post-exposure (trial bins 32-36) blocks. Clouds represent standard error (SE). Bars in panels B, D, F, H depict mean ± SE, while points represent individuals subjects for each respective measure at early-EXP (first 30 trials of EXP) and late-EXP (last 30 trials of EXP) for each group.

For IDE in the left hand, which measured feed-forward reaching error, the mixed-design ANOVA revealed significant main effects of group (*F*(3, 56) = 4.82, *p* < 0.01, *η_g_^2^* = 0.17) and block (*F*(1, 56) = 22.71, *p* < 0.0001, *η_g_^2^* = 0.07). Interference increased over the course of exposure, mostly driven by the visuomotor and combined perturbation groups (Figure 5, C & D). The interaction term was also significant (*F*(3, 56) = 6.90, *p* < 0.001, *η_g_^2^* = 0.06). Examination of simple effects showed that group differences were not statistically significant in early-EXP (*F*(3, 56) = 1.98, *p* = 0.13, *η_g_^2^* = 0.10), but achieved significance in late-EXP (*F*(3, 56) = 7.14, *p* < 0.001, *η_g_^2^* = 0.28). In late-EXP, Tukey HSD showed that the visuomotor group had significantly higher IDE than the controls (*p* < 0.001), and marginally greater IDE than the dynamic perturbation group (*p* = 0.08). Additionally, the combined perturbation group had significantly higher IDE than controls (*p* < 0.01). All other contrasts were not significant after correcting for multiple comparisons. Taken together, these results show that interference due to feed-forward control processes was greatest for the visuomotor and combined perturbation conditions, especially at the end of exposure.

For left-hand IEE, measuring end-point error at the end of the initial ballistic movement, the mixed-design ANOVA revealed significant main effects of group (*F*(3, 56) = 5.85, *p* < 0.01, *η_g_^2^* = 0.19) and block (*F*(1, 56) = 19.16, *p* <0.0001, *η_g_^2^* =0.08), with IEE increasing over the course of the exposure block (Figure 5, E & F). The interaction term was also significant (*F*(3, 56) = 5.07, *p* < 0.01, *η_g_^2^* = 0.06). Within each block, main effects of group were not significant in early-EXP (*F*(3, 56) = 1.39, *p* = 0.25, *η_g_^2^* = 0.07), but achieved significance in late-EXP (*F*(3, 56) = 9.35, *p* < 0.0001, *η_g_^2^* = 0.33). In late-EXP, the visuomotor perturbation (*p* <0.001) and combined perturbation (*p* < 0.01) groups developed significantly higher IEE in their left hand than controls, while in the dynamic group was it marginally different from the controls (*p* = 0.07). Differences between the perturbation groups were not significant after correcting for multiple comparisons. This shows that, at the end of the initial ballistic movement, the visuomotor and combined perturbation groups produced the most interference, particularly at the end of the exposure period, while interference due to the dynamic perturbation of the right hand was lower.

For FEE, measuring the lateral displacement of the left hand from the target once all movement had ceased and incorporating feedback driven corrections at the movement endpoint, the mixed-design ANOVA revealed main effects of group (*F*(3, 56) = 5.08, *p* < 0.01, *η_g_^2^* = 0.14) and block (*F*(1, 56) = 6.27, *p* = 0.02, *η_g_^2^* = 0.04), with FEE decreasing across exposure (Figure 5, G & H). The interaction term was not significant. In early-EXP, the main effect of group was significant (*F*(3, 56) = 3.15, *p* = 0.03, *η_g_^2^* = 0.14), and remained significant at late-EXP (*F*(3, 56) = 3.46, *p* = 0.02, *η_g_^2^* = 0.16). Tukey HSD revealed that that the combined perturbation group had greater deviation than the control group in early-EXP (*p* = 0.06) and in late-EXP (*p* = 0.01). These results show that at the end of movement, the combined perturbation group exhibited the greatest amount of interference, but not over and above that of the other two perturbation groups.

### Kinetic measures

A virtual force channel interspersed pseudorandomly in 20% of trials constrained movement of the left hand to a straight line between home position and targets. Across participants and trials, the average point of peak velocity was at 23% of the whole movement, while the average point of the end of the initial ballistic movement was at 69% of the movement. As such, lateral force applied against the channel wall by the left hand was evaluated between groups at early- and late-EXP at these points, as well as at the end of the movement.

At average peak velocity, early on in the reach, the mixed-design ANOVA revealed significant main effects of both group (*F*(3, 56) = 4.55, *p* < 0.01, *η_g_^2^* = 0.15) and block (*F*(1, 56) = 25.58, *p* < 0.0001, *η_g_^2^* = 0.12), with the lateral force increasing from early-to late-EXP; the interaction term was not significant. Examination of the simple effects in each phase showed a marginal difference between groups in early-EXP: *F*(3, 56) = 2.38, *p* = 0.08, *η_g_^2^* = 0.11) and a significant group difference in late-EXP (*F*(3, 56) = 4.22, *p* < 0.01, *η_g_^2^* = 0.18). In early- and late-EXP, post-hoc tests showed that the visuomotor perturbation group produced greater lateral force against the channel walls than controls (Early-EXP: *p* =0.06; Late-EXP: *p* < 0.01). No other contrasts were significant.

At the end of the initial ballistic movement, ANOVA revealed significant main effects of group (*F*(3, 56) = 5.10, *p* < 0.01, *η_g_^2^* = 0.19) and block (*F*(1, 56) = 8.19, *p* < 0.01, *η_g_^2^* = 0.02), driven by a decrease in lateral force generated by the visuomotor and combined perturbation groups across the exposure period. The interaction term was also significant (*F*(3, 56) = 5.53, *p* < 0.01, *η_g_^2^* = 0.05). Examination of the simple effects showed a significant group main effect at early-EXP (*F*(3, 56) = 7.17, *p* < 0.001, *η_g_^2^* = 0.28), with the combined perturbation (*p* < 0.01) and visuomotor perturbation (*p* < 0.01) generating greater lateral force than the control group. At late-EXP, the group effect was not significant, though visual inspection of the force produced by the left hand showed the dynamic, visuomotor, and combined perturbation groups each producing respectively more force against the channel (Figure 6).

**Figure 6:**
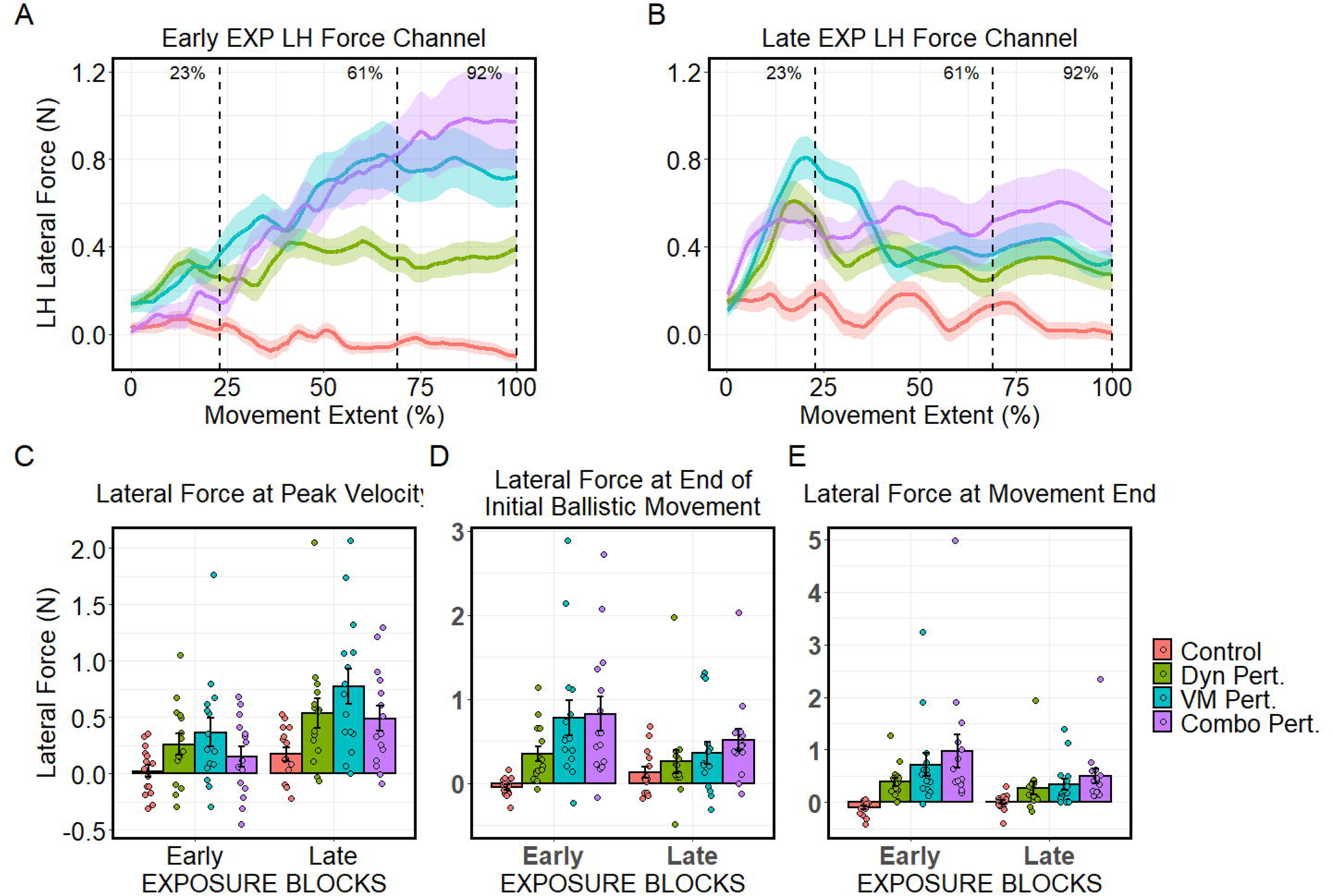
Left hand kinetic results during force channel trials. Top row shows the force applied against the force channel throughout the movement in early-EXP (left panel), and late-EXP (right panel). Bars in C, D, and E depict group mean±SE, while points represent individual subjects at the average point of peak velocity (C), end of the initial ballistic movement (D), and at the end of the movement (E), corresponding 23%, 69%, and 100% of the way through the movement, respectively. Please note that y-axis of C, D, and E were allowed to vary across plots to depict all datapoints.

At the end of the movement, ANOVA showed significant main effects of group (*F*(3, 56) = 5.19, *p* < 0.01, *η_g_^2^* = 0.19) and block (*F*(1, 56) = 11.95, *p* < 0.01, *η_g_^2^* = 0.03), and a significant interaction (*F*(3, 56) = 4.46, *p* < 0.01, *η_g_^2^* = 0.04), again driven by decreasing lateral force generated by the combined and visuomotor perturbation groups across exposure. Examination of simple effects showed significant group differences in early-EXP (*F*(3, 56) = 5.61, *p* < 0.01, *η_g_^2^* = 0.23) and late-EXP (*F*(3, 56) = 3.45, *p* = 0.02, *η_g_^2^* = 0.16). The combined perturbation group generated significantly more force than controls in both early-EXP (*p* < 0.01) and late-EXP (*p* = 0.01), while the visuomotor group generated greater force than controls in early-EXP only (*p* = 0.02).

Taken together, these results show that before participants adapted, the visuomotor and combined perturbation groups produced the greatest amount of force against the channel, particularly at the end of the movement. At the end of the exposure period, once participants had adapted, the visuomotor perturbation group produced the greatest amount of force against the channel early in the movement, while the combined perturbation group produced the greatest force at the end of the movement.

## Discussion

This study examined how adaptation of one hand to different sensorimotor perturbations affects the contralateral hand, manifesting as interference. Importantly, these perturbations targeted different types of sensorimotor information, probing either visuomotor or dynamic processes. First, based on prior experimental data (Desrochers et al., 2017) we predicted that interference in the left hand would be greater when the right hand adapted to a visuomotor perturbation rather than a dynamic perturbation. Overall, this was confirmed in our results; across several dependent measures, visuomotor perturbation resulted consistently in greater interference than dynamic perturbation. This was particularly pronounced in measures that evaluated feed-forward and early feedback driven control (IDE and IEE, respectively), and later in the exposure phase, when participants’ right hand had adapted to the perturbations. Of note, this effect was less pronounced in variables that incorporated feedback driven processes (FEE and RMSE). This may reflect he dominance of visual sensory information in controlling movement trajectories to visual targets (Scheidt *et al*., 2005; Judkins & Scheidt, 2013; Sexton *et al*., 2019).

Further, we hypothesized that interference would be greater when participants were exposed to both perturbations simultaneously. This hypothesis was not supported by our results. In measures of feed-forward driven processes, which assessed movement parameters prior to the integration of reaching feedback (IDE and, to a lesser extent, IEE), the group that received both perturbations in the right hand tended to show roughly equivalent (and sometimes, even slightly less) interference in the left hand compared to the group that received the visuomotor perturbation alone. This occurred despite the combined perturbation being (at peak velocity) larger in magnitude than either the visuomotor or dynamic perturbation. Interestingly, however, in measures that assessed feedback driven processes such as FEE, it was the combined perturbation group that showed the greatest interference. This was also evident in the overall interference measure of movement straightness (RMSE), which likely captured interference during late corrective movements. This pattern of results was also evident from the kinetic measures – at the end of the exposure period, early on in a given movement (i.e., at peak velocity), it was the visuomotor perturbation that showed the greatest interference, and late in the movement (i.e., at the end of the initial ballistic movement and end of the reach), it was the combined perturbation that resulted in greater interference. It is possible that the relatively linear increase with displacement in the force channel kinetics (as shown in Figure 6) may be associated with early learning and/or explicit processes, or changes in limb impedance due to coactivation of muscles in response to the new perturbation. Meanwhile, in late adaptation the force profiles may reflect an updated internal model with modified feed-forward reaching components. Notably, however, the interference seen in the measures incorporating feedback processes (i.e., RMSE, FEE) was not vastly different from that seen for the dynamic and visuomotor perturbations.

These findings are intriguing given previous findings that the sensorimotor system can selectively upregulate its sensitivity to certain types of feedback during learning. For example, participants respond more strongly to a rapid visuomotor perturbation when they are adapting to a dynamic perturbation, as opposed to when they are performing normal reaches (Franklin *et al*., 2012). These responses can also be selectively tuned to different factors affecting the reach, showing that the nervous system can not only learn a specific perturbation, but also modulate secondary mechanisms to adapt to the environment at large (Dimitriou *et al*., 2011; Franklin *et al*., 2014, 2017). Gain-scaling of visuomotor responses also coincides with selective scaling of long-latency stretch reflexes (35-105 ms) during voluntary movements in the early phases of movement learning in unpredictable environments (Pruszynski *et al*., 2011; Cluff & Scott, 2013), suggesting that the nervous system can modulate both feed-forward and feedback control processes during learning. Selectively upregulating sensitivity to certain types of sensorimotor information is thought to allow the sensorimotor system to rapidly correct for further perturbation during environmental uncertainty. Such uncertainty can arise from internal sources such as feedback delays or increased sensorimotor noise (Churchland *et al*., 2006; Faisal *et al*., 2008; van Beers, 2009), or from external sources such as limited feedback or an unpredictable environment (Cluff & Scott, 2013; Nashed *et al*., 2014). Importantly, the ability to scale responses in a context-dependent fashion is a key prediction of optimal feedback control theory (OFTC; (Todorov & Jordan, 2002; Scott, 2004; Todorov, 2005; Diedrichsen *et al*., 2010; Scott *et al*., 2015; Franklin *et al*., 2017)

Despite this compelling evidence of gain scaling in unimanual studies, this study suggests that sensorimotor upregulation may not be shared to the contralateral hemisphere-hand system in our task. If gain scaling parameters are shared via neural crosstalk, the presence of a dynamic perturbation with a visuomotor perturbation in the right hand should have amplified the effects of interference across effectors. Interestingly, research suggests feedback gain responses can be independently controlled by the sensorimotor system during bimanual movements. Brouwer and colleagues (2017) examined corrective responses when cursors were shifted in one hand or in both hands during bimanual reaches and compared resulting motor responses to cursor shifts in unimanual reaches. They found a small but significant decrease in feedback gains during bimanual movements, but also found that the sensorimotor system could independently control responses in each hand. Furthermore, by asymmetrically manipulating the size of the targets to which participants reached, they found that gain responses could independently change within each effector in response to the corresponding target size. As such, each hand could modify its gain response with respect to its own goal. This response is in line with OFCT, which predicts that the sensorimotor system will constrain variability and motor responses if they affect the overall task goal (Diedrichsen, 2007).

In the current experiment, interference in the combined perturbation group may not be increased beyond that of the visuomotor perturbation alone because the left hand does not receive forces with similar characteristics of those in the right hand. In other words, since the left hand is not required to interact with a dynamic perturbation, that limb will not be susceptible to the effects of the perturbation in the right hand. This may be due to dynamic adaptation occurring in independent, intrinsic frames of reference specific to the adapting hand (Shadmehr & Mussa-Ivaldi, 1994). Thus, these results suggest that feedback gains within each hemisphere-hand system can be independently regulated in parallel with one another. Further, it suggests that these upregulated feedback-gains in one hemisphere-hand system are not communicated between hemispheres via neural crosstalk.

This study also raises interesting questions as to why visuomotor perturbations produce greater interference than dynamic perturbations during bimanual movements. One could argue that a simple answer may lie in the fact that in the visuomotor perturbation task, the movements of the hands are inherently asymmetrical, as the right hand must change its trajectory to hit the target. However, the time course of interference in our task, which increases gradually over the course of exposure, lends support to interference being enhanced as a result of adaptive processes – if interference was strictly due to asymmetrical reaches alone, it would be present at its maximal magnitude as soon as the task required the participants to reach asymmetrically. Additionally, most participants anecdotally reported that they were unaware that their reaches were asymmetrical by the end of the exposure period in the visuomotor condition. Instead, they perceived their movements as being symmetrical – yet interference remained present towards the end of exposure. Thus, the differences in interference between directional and adaptation-driven perturbation types is likely, at least in part, driven by neural processes tied to adapting to the visuomotor perturbation itself.

As such, the neural circuitry involved in adapting to these perturbations may play an integral role in the observed differences in interference. Research suggests that adaptation to visuomotor and dynamic perturbations may recruit slightly different neural substrates, particularly in the cerebellum (Rabe *et al*., 2009; Donchin *et al*., 2011). Additionally, visuomotor control also requires a complex premotor-parietal network responsible for localizing targets in space and planning movement trajectories, in addition to contribution of M1 (Kurata, 1994; Desmurget *et al*., 1999; Tanaka *et al*., 2009; Mutha *et al*., 2011; Werner *et al*., 2014; Culham, 2015; Manuweera *et al*., 2018). Critically, visuospatial information is also projected to both hemispheres, requiring posterior parietal cortices of both hemispheres to successfully plan spatial movements (Goodale & Milner, 1992). As such, is not surprising that severing portions of the corpus callosum that connect posterior parietal cortex preferentially abolishes spatial interference (Eliassen *et al*., 1999). Further, the left parietal lobe is particularly involved in visuomotor adaptation (Mutha *et al*., 2011); this may contribute to the elevated role of visuomotor feedback on interference demonstrated in this study. It is also possible that visuomotor processes hold a privileged status in being shared between hemispheres via the corpus callosum over dynamic processes. However, whether each type of adaptation elicits differential patterns of activity are equivocal, and dynamic adaptation may share several common brain areas with visuomotor perturbations, including some parietal and cerebellar loci (Krakauer *et al*., 2004; Diedrichsen *et al*., 2005; Graydon *et al*., 2005; Ferrari-Toniolo *et al*., 2015). Additionally, neural responses to different perturbations may change depending on how engaged brain regions functionally interact throughout the acquisition of adaptive responses (Tunik *et al*., 2007). Furthermore, while interhemispheric contributions bimanual coordination and interference are well established (Franz *et al*., 1991; Andres *et al*., 1999; Gerloff & Andres, 2002; Swinnen, 2002; Hazeltine *et al*., 2003; Desrochers *et al*., 2020), recent evidence has demonstrated that neuronal activity in M1 of both hemispheres are functionally related to the action of a single arm (Ames & Churchland, 2019; Heming *et al*., 2019). Thus, it is possible that interference may also result from ipsilateral contributions of the left hemisphere. Future work could attempt to further understand ipsilateral M1 contributions to bimanual movements (Cross *et al*., 2020), the contribution of ipsilateral vs. contralateral activity in the context of interference, and whether these processes interact with different types of sensorimotor information.

These perturbations are also functionally distinct. Visuomotor and dynamic perturbations do not interfere with one another (Krakauer *et al*., 1999; Tcheang *et al*., 2007), and likely use different reference frames (Shadmehr & Mussa-Ivaldi, 1994; Flanagan & Rao, 1995). However, more recent evidence suggests that multiple reference frames may be utilized depending on the context of the movement (Berniker *et al*., 2013; Parmar *et al*., 2015) and that perturbations do indeed interact when applied to the same motor variable (Tong *et al*., 2002). As such, it is possible that visuomotor perturbations affect reference frames that are shared between effectors, whereas dynamic perturbations utilize reference frames that are isolated in a single effector. If true, then manipulating reference frames to become more shared during dynamic perturbations could increase interference.

Previous findings also point to a role of explicit learning during adaptation to visuomotor rotations (Taylor *et al*., 2014; Bond & Taylor, 2015). While interference is most likely an implicit phenomenon, occurring in the left hand that moves without visual feedback, an open question remains as to whether interference is itself driven by implicit or explicit adaptation processes in the right hand. It could be that explicit components of visuomotor adaptation deliver a greater neural drive to intermanual interference than do implicit processes (i.e., updating of internal models and gain scaling), which could account for the pattern of results observed in this study. At the same time, it should be noted that the concurrent bimanual activity, with one hand adapting and the other ‘flying blind’, is certainly more difficult to perform than a unimanual task. Participants’ attention is fully occupied in simultaneously moving to two targets, and it is plausible to assume that there are fewer resources available for forming explicit strategies while performing the demanding task. Interestingly, recent evidence also suggests that explicit learning is at play during adaptation to dynamic perturbations as well (Schween *et al*., 2020). Thus, explicit processes alone may not fully explain the observed differences in interference between the visuomotor and dynamic perturbation groups in the current study. This motivates further study as to whether implicit or explicit processes of adaptation in one hand differently affect interference in the opposing hand.

This work has key implications not only for understanding bimanual control in healthy individuals, but also for those with impaired movement. Understanding the phenomenon of interference may help to elucidate mechanisms behind “mirror movements”, a symptom that can occur in some movement disorders including Parkinson’s disease and dystonia (Cox *et al*., 2012). Additionally, bimanual coordination training is a potential strategy to facilitate recovery of function in hemiplegic upper limbs in stroke patients (Rose & Winstein, 2004; Sainburg *et al*., 2013). Understanding how sensorimotor information is shared between hemispheres may inform or improve these approaches.

This study is not without some limitations. First, the combined perturbation group consisted of perturbations that each forced participants to make counterclockwise corrective actions (either reach direction or force). It was anticipated that this would be the best method to elicit the largest amount of interference in left hand. However, previous research has suggested that some individuals may show mirrored patterns of interference, as opposed to interference in the same direction (Kagerer, 2015b). As such, these perturbations may have acted to inhibit the observed interference, each cancelling out left-hand effects from the other. We deem this unlikely, since all participants exhibited interference in the same direction as the perturbations. However, future studies could combine perturbations of different directions, and observe how left-hand interference is modulated as a result of these different directions. This could provide additional information into how visuomotor and dynamic perturbations interact within the sensorimotor system. Second, conditions in which asymmetrical reaches were performed without the presence of a visuomotor rotation were not included. This was because the main objective of the study was to see if concurrent dynamic adaptation upregulated the effects of the visuomotor perturbation. Indeed, new data from our lab indicate that the presence of adaptive processes themselves increase interference above asymmetrical reaching alone (Brunfeldt et al., under review). In this study, a bimanual task similar to the one used in this experiment was contrasted to a task in which participants had to move the right hand under veridical visual feedback to targets that were rotated by 40° while the left hand moved under kinesthetic feedback to targets placed straight forward and behind the home position. The results showed that left-hand interference was smaller in that task than in the task that required adaptation to a 40° visuomotor feedback rotation, and that the time course of the interference was different. This suggests that it is the adaptive component in this task that drives most of the interference. Further, research showing upregulation of visuomotor feedback gain also uses a shifted cursor to probe the visuomotor activity (Franklin *et al*., 2012). Given that the combined perturbation did not show increased interference, it is also unlikely that a dynamic perturbation combined with non-adaptive asymmetrical reaching would have resulted in increased patterns of interference, though future studies will examine this. Third, we used a velocity dependent curl field as our dynamic perturbation, as this type of force field tends to be used more often in the literature. However, a position-dependent force field (e.g., Sing *et al*., 2009) may have allowed for better comparison between the dynamic and visuomotor perturbations, as the magnitude of the perturbation would be better matched over the course of the reach. Nevertheless, if adapting to one type of sensorimotor perturbation upregulates sensorimotor feedback gains and increases sensitivity to a different sensorimotor perturbation (Franklin *et al*., 2012, 2017), the velocity dependent force field utilized in the present study should still have produced upregulation of sensorimotor feedback gains in the combined perturbation group.

In summary, this study has shown that visuomotor and dynamic perturbations with similar magnitude show different levels of left-hand interference, with the visuomotor perturbation eliciting greater interference, particularly in feed-forward measures at the end of an adaptation period. Further, we have shown that simultaneously adapting to visuomotor and dynamic perturbations in one hand does not increase interference in the opposing hand beyond that of a visuomotor perturbation alone. This was demonstrated by similar levels of left-hand interference when participants received visuomotor perturbation or simultaneous visuomotor and dynamic perturbations in the right hand. Future work will determine if this may be the result of upregulated sensorimotor feedback gains not transferring between hemispheres.

## Acknowledgements

The authors would like to thank the individuals who participated in this study. This research was supported by Michigan State College of Education Summer Research Fellowships (to P.C.D. and A.T.B.), the MSU Graduate School Research Enhancement Award (to P.C.D.), and the Dykstra Family Research Endowment in Education (to P.C.D.).

## Author Contributions

P.C.D. conceived of and designed the experiment, with input from A.T.B. and F.A.K. P.C.D. collected the data and performed statistical analyses. Figures were made by P.C.D. The original manuscript draft was written by P.C.D. Revision and editing was provided by P.C.D., A.T.B., and F.A.K.

## Data Accessibility

Data and analysis code can be found in P.C.D.’s GitHub repository at https://github.com/philcd89

## Abbreviations

ANOVA: Analysis of Variance
Combo Pert: Combined Perturbation
DYN Pert: Dynamic Perturbation
EXP: Exposure
FEE: Final Endpoint Error
IDE: Initial Directional Error
IEE: Initial Endpoint Error
KBL: Kinesthetic baseline
OFCT: Optimal Feedback Control Theory
Post-EXP: Post-exposure
RMSE: Root mean square error
VBL: Visual baseline
VM Pert: Visuomotor perturbation

